# Synchronization of human retinal pigment ephitilial-1 (RPE-1) cells in mitosis

**DOI:** 10.1101/2020.04.21.052803

**Authors:** Stacey J. Scott, Kethan Suvarna, Pier Paolo D’Avino

## Abstract

Human retinal pigment ephitilial-1 (RPE-1) cells are increasingly being used as a model to study mitosis because they represent a non-transformed alternative to cancer cell lines, such as HeLa cervical adenocarcinoma cells. However, the lack of an efficient method to synchronize RPE-1 cells in mitosis precludes their application for large-scale biochemical and proteomics assays. Here we report a protocol to synchronize RPE-1 cells based on sequential treatments with the Cdk4/6 inhibitor PD 0332991 (palbociclib) and the microtubule depolymerizing drug nocodazole. With this method, the vast majority (80-90%) of RPE-1 cells arrested at prometaphase and exited mitosis synchronously after release from nocodazole. Furthermore, we show that this protocol could be successfully employed for the characterization of the protein-protein interaction network of the kinetochore protein Ndc80 by immunoprecipitation coupled with mass spectrometry. This synchronization method significantly expands the versatility and applicability of RPE-1 cells to the study of cell division and might be applied to other cell lines that do not respond to treatments with DNA synthesis inhibitors.

## INTRODUCTION

Cell division is essential for growth, development and reproduction in most organisms and errors during this process are responsible or have been implicated in several human diseases, including cancer pathologies. The process of mitosis ensures the faithful and accurate segregation of both genomic and cytoplasmic contents into two daughter cells. Mitosis progresses through a series of phases -prophase, prometaphase, metaphase, anaphase and telophase-during which the microtubule spindle is assembled in order to align and segregate the duplicated chromatids (Reber and Hyman, 2015). The centromeric regions of chromatids attach to spindle microtubules through the kinetochore, a macromolecular structure composed of a multitude of proteins and protein complexes (Cheeseman, 2014). The kinetochore is divided into two layers, the inner and outer kinetochore. The inner kinetochore comprises many CENP proteins that assemble to form the constitutive centromere-associated network (CCAN) (Cheeseman, 2014) while the outer kinetochore is comprised primarily of the large multi-subunit Knl1/Mis12/Ndc80 complex network (KMN network), which is recruited by the CCAN at the inner kinetochore to form strong interactions with mitotic spindle microtubules (Cheeseman et al., 2006). After all chromatids form correct bipolar attachments, cyclin B is degraded and cells can exit mitosis and segregate chromatids towards opposite poles during anaphase (Wieser and Pines, 2015). After anaphase onset, the mitotic spindle is reorganized into an array of antiparallel and interdigitating microtubules known as the central spindle that, together with the contraction of an equatorial actomyosin ring, drives the separation of the two daughter cells during cytokinesis, completing the cell division process (D’Avino et al., 2015).

A combination of genetics, biochemical, and high-resolution imaging techniques is required to finely dissect the mechanics and regulation of mitotic events (Ong and Torres, 2019). Cultured human cells have proven a very useful model for the study of mitosis, but most of the cell lines employed in the field originated from tumors and thus often have alterations in the main genes and signaling pathways that regulate cell growth, cell cycle, and mitosis. For example, HeLa cervical adenocarcinoma cells (Jones et al., 1971) are widely used in mitotic studies because they are easy to image, grow in large quantities, and can be easily synchronized at different mitotic stages using a thymidine-nocodazole block and release method. However, HeLa cells are transformed by the human papilloma virus, which silences the crucial tumor suppressor p53 (Scheffner et al., 1991; Yee et al., 1985). Moreover, these cells are genetically unstable and near-tetraploid, which makes the use of gene-editing techniques laborious and time-consuming because of the presence of multiple gene copies. For these reasons, the diploid, non-transformed, hTERT-immortalized, retinal pigment ephitelial-1 (RPE-1) cell line (Bodnar et al., 1998) is emerging as a very valid alternative to HeLa for the study of mitosis. However, an effective method to synchronize these cells in mitosis is still lacking, which precludes their use for large-scale biochemical and molecular studies aimed at dissecting the functions, regulation, and interactions of mitotic proteins. Here we describe a protocol, based on combined sequential treatments with the Cdk4/6 inhibitor PD 0332991 (palbociclib) (Fry et al., 2004) and the microtubule depolymerizing drug nocodazole (Sentein, 1977), that can be used to efficiently synchronize RPE-1 cells in mitosis. We also demonstrate that this synchronization method can be successfully employed for the characterization of the protein-protein interaction network (interactome) of the kinetochore protein Ndc80 by immunoprecipitation coupled with mass spectrometry.

## RESULTS AND DISCUSSION

### Synchronization of RPE-1 cells using a palbociclib-nocodazole block and release method

Our initial attempts to synchronize RPE-1 and other p53 wild-type cells in mitosis using DNA synthesis inhibitors, like thymidine or aphidicolin, failed because these drugs presumably triggered a DNA damage response and consequent p53 activation and cell cycle arrest (data not shown). To overcome this problem, we decided to employ the Cdk4/6 inhibitor palbociclib to instead block cells in the G1 phase of the cell cycle (Fry et al., 2004). RPE-1 cells were treated with palbociclib for 18 hours, released for 8 hours and then incubated with nocodazole for 12 hours. After nocodazole treatment, cells were extensively washed, released in fresh medium, and collected at different time intervals (Fig. 1A; see also Materials and Methods). We observed almost no round-up mitotic cells after palbociclib treatment, indicating that the cells arrested in G1 (Fig. 1B). By contrast, the vast majority of cells (80%-90%) after incubation with nocodazole appeared to be in mitosis (Fig. 1B).

**Fig. 1.**
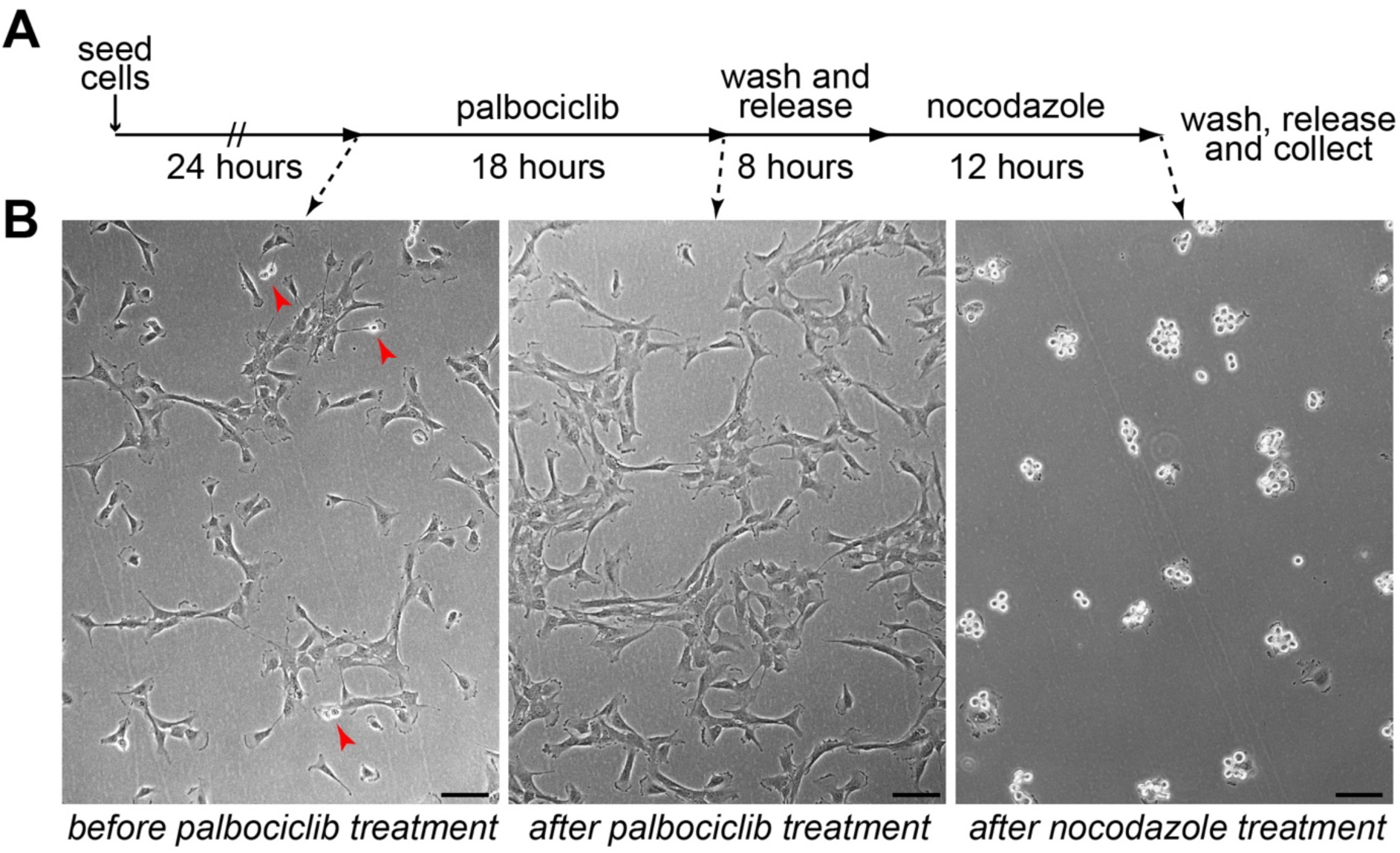
RPE-1 cells can be synchronized in mitosis using a palbociclib-nocodazole block. (A) Schematic diagram of the synchronization protocol. (B) Microscopy images of RPE-1 cells before and after palbociclib treatment (left and middle panels), and after nocodazole incubation (right panel). The red arrowheads in the left panel mark dividing cells. Note the vast majority of round dividing cells after nocodazole treatment. Bars, 100 µm.

We next analyzed RPE-1 cells collected at 0, 50, and 75 min after nocodazole release by both Western blot and flow cytometry analysis to identify their mitotic stages (Fig. 2). We also included a sample of cells released for 50 min in medium containing the proteasome inhibitor MG132, which should prevent cyclin B degradation and mitotic exit. The levels of both cyclin B and the mitotic marker histone H3 pS10 dropped by more than 70-80% after 50 min (Fig. 2A), indicating that most cells had exited mitosis. By contrast, the levels of both mitotic markers were unaltered or even increased in cells released for 50 min in medium with MG132 (Fig. 2A). Flow cytometry profiles indicated that more than 90% of the cells at the 0 min time-point had 4*n* DNA content and that this percentage gradually decreased after 50 and 75 min. This decrease in 4*n* cells was paralleled by an increase in the number of 2*n* G1 cells, which were the majority in the 75 min sample (Fig. 2B). We further refined our Western blot analysis in samples collected at more frequent intervals (30, 45, 60, and 75 min) after nocodazole release (Fig. 2C). Cyclin B levels dropped by 70% after just 30 min, while a much more modest decrease of less than 40% was observed for histone H3 pS10 (Fig. 2C). By contrast, the levels of both markers only marginally decreased in cells released for 30 min in the presence of MG132 (Fig. 2C). Cyclin B and histone H3 pS10 levels gradually decreased in the 45, 60, and 75 min samples to 20% or less compared to the 0 time point (Fig. 2C).

**Fig. 2.**
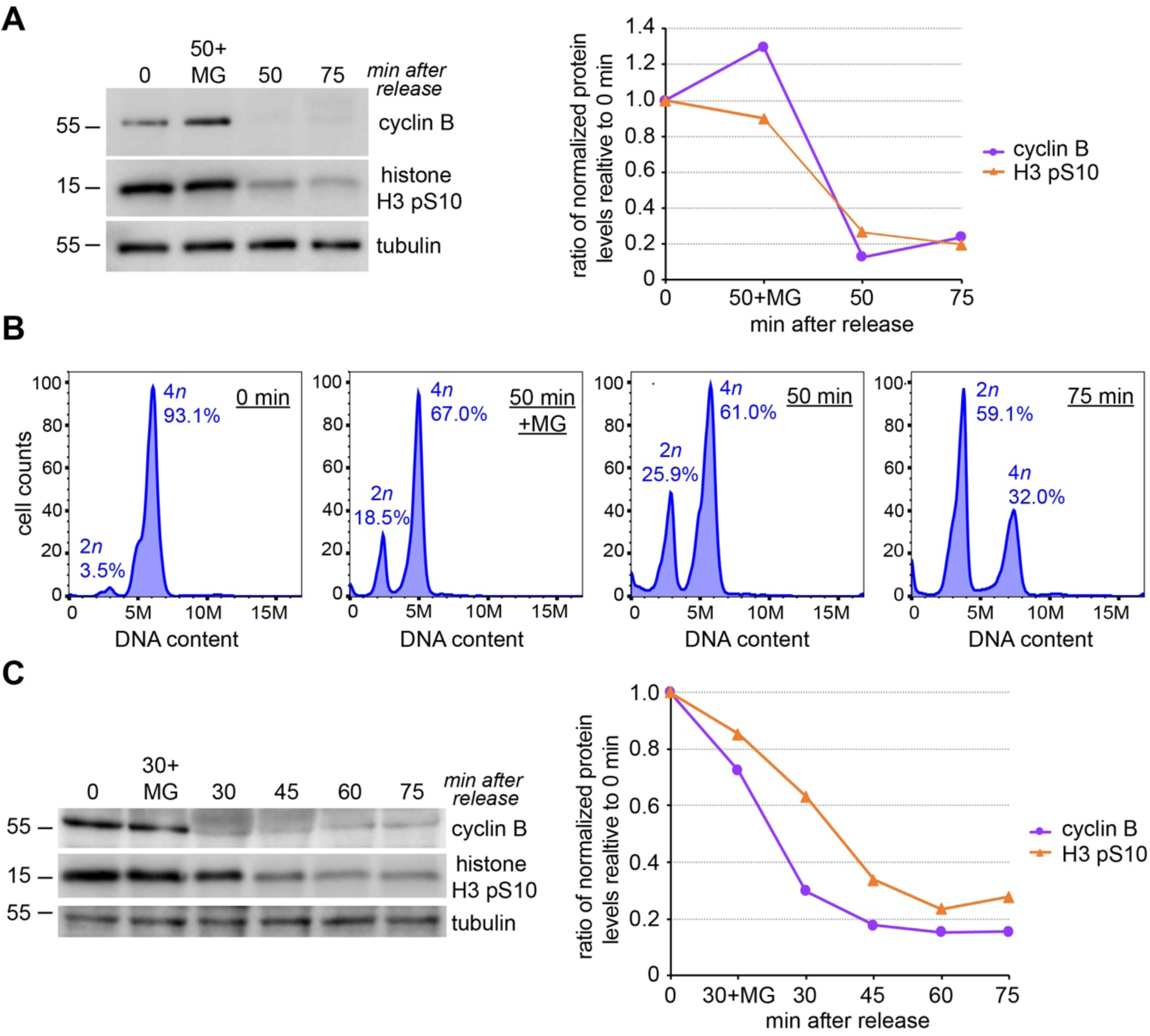
RPE-1 cells exit mitosis synchronously after release from the nocodazole block. (A) Analysis of protein expression in RPE-1 cells released from nocodazole. Proteins were extracted from RPE-1 cells at the times indicate at the top, separated by SDS PAGE, and analyzed by Western blot to identify the proteins indicated to the right. The numbers on the left indicate the sizes of the molecular mass marker. The graph at the right shows the quantification of protein levels, normalized to tubulin and relative to levels at time 0. (B) Flow cytometry profiles of cells released from the nocodazole block at the same time points as in (A) and stained with propidium iodide (PI) to analyze DNA content. (C) Proteins were extracted from RPE-1 cells at the times indicate at the top, separated by SDS PAGE, and analyzed by Western blot to identify the proteins indicated to the right. The numbers on the left indicate the sizes of the molecular mass marker. The graph at the right shows the quantification of protein levels, normalized to tubulin and relative to levels at time 0.

Together, these results indicate that most RPE-1 cells are in anaphase 30 min after release from nocodazole and that the majority of cells have completed mitosis and are in G1 after 60-75 min.

### Characterization of the Ndc80 interactome using the palbociclib-nocodazole block and release method

To demonstrate the applicability of our synchronization protocol, we decided to characterize the interactome of the outer kinetochore component Ndc80 in RPE-1 cells synchronized at metaphase. We used our palbociclib-nocodazole block and release method to synchronize and collect RPE-1 cells 30 min after release from nocodazole in medium containing MG132. Protein extracts from these cells were used in immuno-precipitation (IP) experiments with antibodies against the outer kinetochore component Ndc80 or general mouse IgG as control and then baits and prey proteins were identified by mass spectrometry (Fig. 3). After eliminating non-specific preys (see Materials and Methods), we identified 380 Ndc80-specifc interactors (Table S1), which included all the four components of the Ndc80 complex (Ndc80, Nuf2, Spc24 and Spc25) as well as many other proteins known to be involved in spindle assembly and chromosome attachment and alignment (Tables 1 and S1). As expected, the Gene Ontology (GO) enrichment profile of the Ndc80 interactome indicated a strong enrichment in proteins involved in mitosis, cell cycle, spindle assembly and microtubule dynamics (Fig. 3B, Table S2). However, we also unexpectedly found a significant enrichment in proteins involved in RNA processing and splicing (Fig. 3B, Supplementary Data 2). This finding suggests a potential role for these proteins, and possibly RNA, in mitosis that deserves future investigation.

**Fig. 3.**
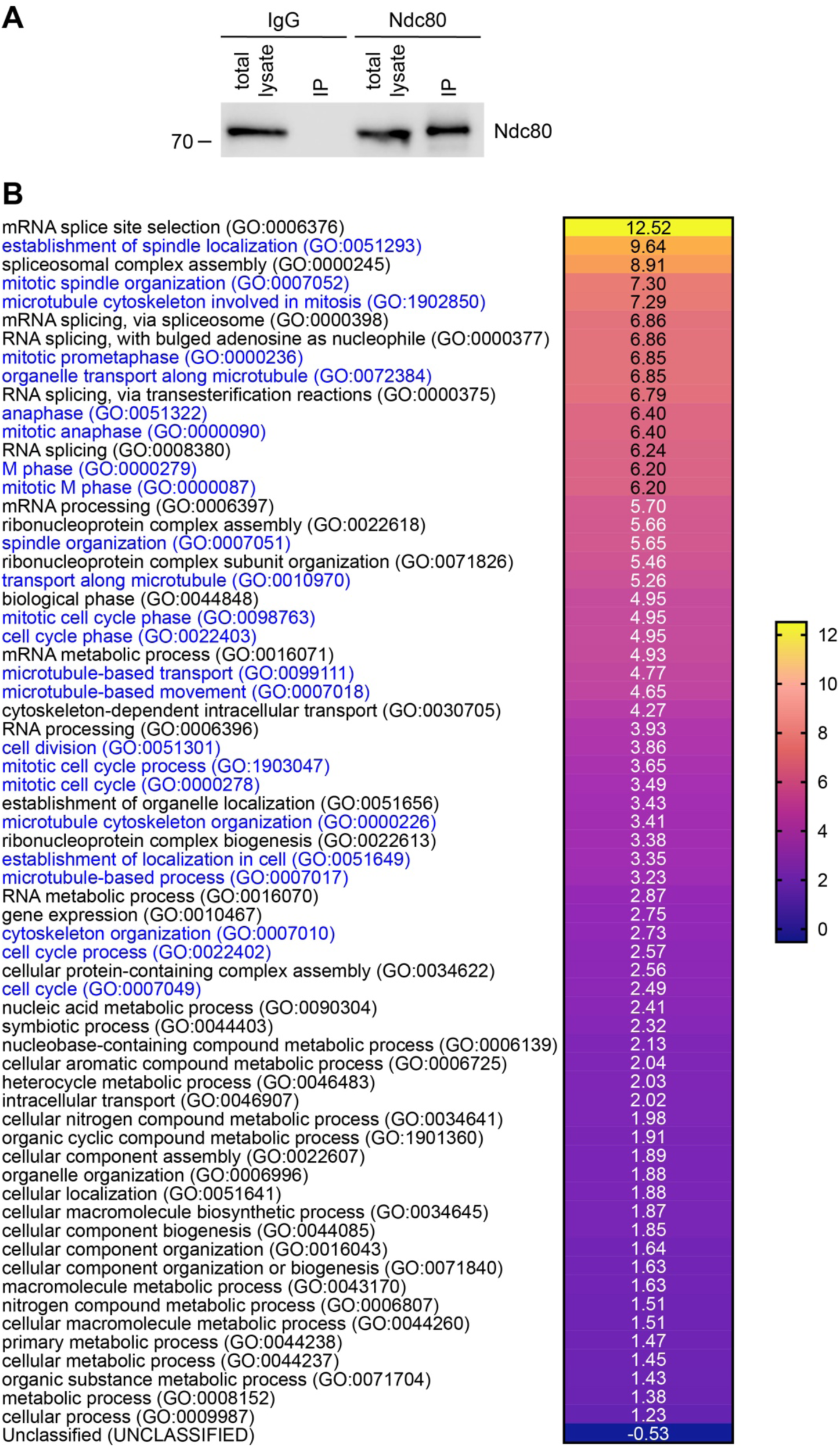
Characterization of the Ndc80 interactome in metaphase RPE-1 cells. (A) Protein extracts were used in IP assays using either Ndc80 or IgG and then extracts and pull-downs were analyzed by Western blot to detect Ndc80. The numbers on the left indicate the sizes of the molecular mass marker. (B) Heat map showing the GO annotation enrichment profile of the Ndc80 interactome analyzed using PANTHER under the category GO biological process. Overrepresented GO terms are shown in different color shades according to their fold enrichment as indicated in the color scale bar at the right; actual fold enrichment values are shown within the heat map. GO terms for processes involved in cell cycle, mitosis and microtubule dynamics are in blue. Only results for Bonferroni-corrected for *p* <0.05 were considered (see Table S2).

**Table 1.** List of selected proteins involved in chromosome attachment and alignment identified in the Ndc80 interactome.

| **Name** | **Function** | **Score** | **Peptides** |
| --- | --- | --- | --- |
| **NDC80** | Kinetochore protein, Ndc80 complex subunit | 1462 | 42 |
| **NUF2** | Kinetochore protein, Ndc80 complex subunit | 867 | 24 |
| **SPC24** | Kinetochore protein, Ndc80 complex subunit | 765 | 33 |
| **SPC25** | Kinetochore protein, Ndc80 complex subunit | 399 | 11 |
| **KNL1** | Kinetochore protein | 53 | 1 |
| **NSL1** | Kinetochore protein, Mis12 complex subunit | 33 | 1 |
| **ZWINT** | ZW10 interactor | 181 | 3 |
| **CENP-E** | Centromere-associated protein E | 51 | 1 |
| **INCENP** | Inner centromere protein, CPC component | 65 | 1 |
| **PP1A** | Serine/threonine phosphatase PP1α catalytic subunit | 2222 | 49 |
| **PP1B** | Serine/threonine phosphatase PP1β catalytic subunit | 2033 | 43 |
| **PP1G** | Serine/threonine phosphatase PP1γ catalytic subunit | 2127 | 47 |
| **PP2AA** | Serine/threonine phosphatase 2A catalytic subunit $\alpha$ isoform | 49 | 2 |
| **PLK1** | Serine/threonine-protein kinase Polo-like kinase 1 | 102 | 2 |
| **DYHC1** | Cytoplasmic dynein 1 heavy chain 1 | 1500 | 55 |
| **DC1L1** | Cytoplasmic dynein 1 light intermediate chain 1 | 84 | 1 |
| **DC1L2** | Cytoplasmic dynein 1 light intermediate chain 2 | 47 | 2 |
| **DCTN1** | Dynactin subunit 1 | 34 | 1 |

Protein names are according to the UniProt database. The Mascot score (Score) and number of peptides identified are indicated. Proteins are listed and grouped by function rather than score and peptide numbers, with members of the Ndc80 complex listed at the top.

### Concluding remarks

Our results demonstrate that combined and sequential treatments with the Cdk4/6 inhibitor palbociclib and the microtubule depolymerizing drug nocodazole can be successfully used to synchronize RPE-1 cells at different mitotic stages. This method allows the large-scale purification of RPE-1 synchronized cells that can be used for biochemical and proteomics studies, including the identification of protein networks and characterization of post-translational modifications of the mitotic proteome. We believe that this new technique will significantly expand the versatility and applicability of RPE-1 cells to the study of cell division. Finally, our preliminary results indicate that this protocol can be applied to other cell lines that have a functional p53 response or do not respond to treatments with DNA synthesis inhibitors (data not shown).

## MATERIALS AND METHODS

### Cell culture, synchronization and imaging

hTERT-RPE-1 cells (ATCC) were cultured in DMEM F-12 + Glutamax medium (GIBCO) supplemented with 10% fetal bovine serum (Sigma-Aldrich), 1% penicillin-streptomycin (ThermoFisher), and 0.01 mg ml^-1^ hygromycin B (ThermoFisher) at 37°C and 5% CO_2_.

For synchronization, RPE-1 cells were seeded at 1/6 confluence. After 24 hours, palbociclib (PD 0332991, Selleckchem) was added to the medium at a final concentration of 1 µM and cells incubated for further 18 hours. After three washes with phosphate-buffered saline (PBS + CaCl_2_ and MgCl_2_) to remove the drug, cells were cultured for 8 hours in fresh complete medium before adding 50 ng ml−1 nocodazole (Sigma-Aldrich). After 12 hours, cells were harvested by mitotic shake-off, centrifuged at 1000 *g* for 3 min, washed five times with large volumes of PBS + CaCl_2_ and MgCl_2_, and released in fresh complete medium with or without 10 μM MG132 (Sigma-Aldrich) for the appropriate times before collection.

Cells were imaged using a Leica DMi1 inverted microscope with integrated camera.

### Western blot analysis

Cells were centrifuged, resuspened in phosphate buffer saline (PBS) and then an equal volume of 2x Laemmli buffer was added. Samples were then boiled for 10 min and stored at -20°C. Proteins were separated by SDS PAGE and then transferred onto PVDF membrane (Immobilon-P) at 15V for 1 hour. Membranes were blocked overnight at 4°C in PBS + 0.1% (v/v) Tween (PBST) with 5% (w/v) dry milk powder. After blocking, membranes were washed once with PBST and then incubated with the appropriate primary antibody diluted in PBST + 3% (w/v) BSA (Sigma) for 2 hours at RT. Membranes were washed 3×5 minutes in PBST and then incubated with HRP-conjugated secondary antibodies in PBST + 1% BSA for 1 hour at room temperature. After further 3×5 min washes in PBST, the signals were detected using the ECL West Pico substrate (ThermoFisher) and chemiluminescent signals were acquired below saturation levels using a G:BOX Chemi XRQ (Syngene) and quantified using Fiji (Schindelin et al., 2012).

### Flow cytometry

RPE-1 cells were centrifuged at 1000 *g* for 3 min, the supernatant removed, and cell pellets resuspended in 5 ml of PBS. 2 ml of ice-cold 70% ethanol was added drop-wise to the pellet while vortexing and cells maintained at -20°C for a minimum of 2 hours. After centrifuging at 800 *g* for 5 minutes at 4°C, the supernatant was removed and cell pellets were washed twice with PBS before resuspending in 0.2-0.5 ml of propidium iodide (PI)/RNAse staining buffer (BD Pharmigen) followed by incubation at room temperature for 15 minutes in the dark. Stained cells were then stored at 4°C for a minimum of 24 hours, strained through a 35 µm nylon mesh, and loaded onto a CytoFLEX S (Beckman Coulter) instrument. Cytexpert software (Beckman Coulter) was used for data acquisition, a 561nm laser with a 610/20 filter was used for detection, and 10000 single cell events were recorded for each sample. Annotation of data was performed manually using FlowJo software.

### Immunoprecipitation (IP)

100 µl of Dynabeads Protein A (ThermoFisher) were prepared for immunoprecipitation (IP) by washing 3 times with phosphate-buffer saline (PBS) followed by incubation with either 10 µg of mouse monoclonal anti-Ndc80/Hec1 antibody (Santa Cruz, sc-515550) or 10 µg of mouse IgG (Sigma-Aldrich) in 200 µl of PBS + 0.1% (v/v) NP40 on a rotating wheel at 4°C overnight.

Six 175 cm^2^ flasks of RPE-1 cells were seeded and synchronized as described above. Cells were collected after 30 min incubation in fresh medium containing 10 μM MG132 (Sigma-Aldrich), washed, centrifuged and the cell pellet stored at -80°C. Cell pellets were thawed by directly resuspending in 2 ml of extraction buffer (EB: 50 mM HEPES pH 7, 100 mM KAc, 50 mM KCl, 2 mM MgCl_2_, 1 mM EGTA, 0.1% [v/v] NP-40, 1 mM DTT, 5% [v/v] glycerol and Roche Complete Protease Inhibitors) and homogenized using a high-performance disperser (Fisher). The cell lysate was treated with 200 mg/ml of DNAse I (New England BioLabs) for 10 min at 37°C followed by 10 min at room temperature and then clarified by centrifugation at 1500 *g* for 15 min at 4°C. After centrifugation, the supernatant was split into two and each sample was added to the Dynabeads Protein A pre-incubated with either the Ndc80 antibody or mouse IgG (see above) on a rotating wheel at 4°C for 4 hours. Beads were then washed four times using a magnetic stand in 5 ml of EB for 5 min on a rotating wheel at 4°C, transferred to a new tube, and washed one more time with 5 ml of PBS. After removing as much liquid as possible, beads were stored at -80°C before being analyzed by liquid chromatography coupled with tandem MS (LC-MS/MS; see section below).

#### Mass spectrometry (MS) analyses

For the analysis of IP samples, beads were digested with trypsin and processed as previously described (McKenzie et al., 2016). Raw MS/MS data were analyzed using the MASCOT search engine (Matrix Science). Peptides were searched against the UniProt human sequence database and the following search parameters were employed: enzyme specificity was set to trypsin, a maximum of two missed cleavages were allowed, carbamidomethylation (Cys) was set as a fixed modification, whereas oxidation (Met), phosphorylation (Ser, Thr and Tyr) and ubiquitylation (Lys) were considered as variable modifications. Peptide and MS/MS tolerances were set to 25 parts per million (ppm) and 0.8 daltons (Da). Peptides with MASCOT Score exceeding the threshold value corresponding to <5% False Positive Rate, calculated by MASCOT procedure, and with the MASCOT score above 30 were considered to be positive.

#### Computational and statistical analyses

We used in-house written Perl scripts to compare the Mascot data from the Ndc80 and IgG IP experiments in order to eliminate non-specific hits. Prey hits either absent from or having ≥5-fold increase in both MASCOT score and peptide numbers compared to the IgG negative control were classed as being specific. Additional common contaminants, such as keratins and hemoglobin, were eliminated manually.

GO enrichment analysis was performed using PANTHER (Mi et al., 2017). Prism8 (GraphPad) and Excel (Microsoft) were used for statistical analyses and to prepare graphs.

## Acknowledgments

We thank Luisa Capalbo for critical reading of the manuscript.

## Author contributions

Conceptualization: PPD and SJS; Investigation: SJS and KS; Formal analysis: SJS, KS and PPD; Funding acquisition: PPD; Methodology: SJS, KS and PPD; Project administration: PPD; Supervision: PPD; Validation, SJS, KS and PPD; Visualization: SJS, KS and PPD; Writing – original draft: PPD; Writing – review and editing: SJS, KS and PPD.

---

**Table S1.** Excel file listing the proteins identified by MS from the Ndc80 IP of RPE-1 cells synchronized in metaphase. Non-specific binding proteins were removed by filtering the Ndc80 dataset against the IgG dataset (see Materials and Methods). The second sheet lists the same protein with their respective gene names and GO terms.

**Table S2.** Excel file showing the results of the GO enrichment analyses from the MS data of the Ndc80 IP shown in Table S1.

## References

Bodnar, A. G., Ouellette, M., Frolkis, M., Holt, S. E., Chiu, C. P., Morin, G. B., Harley, C. B., Shay, J. W., Lichtsteiner, S. and Wright, W. E. (1998). Extension of life-span by introduction of telomerase into normal human cells. Science 279, 349–52.

Cheeseman, I. M. (2014). The kinetochore. Cold Spring Harb Perspect Biol 6, a015826.

Cheeseman, I. M., Chappie, J. S., Wilson-Kubalek, E. M. and Desai, A. (2006). The conserved KMN network constitutes the core microtubule-binding site of the kinetochore. Cell 127, 983–97.

D’Avino, P. P., Giansanti, M. G. and Petronczki, M. (2015). Cytokinesis in animal cells. Cold Spring Harb Perspect Biol 7, a015834.

Fry, D. W., Harvey, P. J., Keller, P. R., Elliott, W. L., Meade, M., Trachet, E., Albassam, M., Zheng, X., Leopold, W. R., Pryer, N. K. et al. (2004). Specific inhibition of cyclin-dependent kinase 4/6 by PD 0332991 and associated antitumor activity in human tumor xenografts. Mol Cancer Ther 3, 1427–38.

Jones, H. W., Jr., McKusick, V. A., Harper, P. S. and Wuu, K. D. (1971). George Otto Gey. (1899-1970). The HeLa cell and a reappraisal of its origin. Obstet Gynecol 38, 945–9.

McKenzie, C., Bassi, Z. I., Debski, J., Gottardo, M., Callaini, G., Dadlez, M. and D’Avino, P. P. (2016). Cross-regulation between Aurora B and Citron kinase controls midbody architecture in cytokinesis. Open Biol 6, 160019.

Mi, H., Huang, X., Muruganujan, A., Tang, H., Mills, C., Kang, D. and Thomas, P. D. (2017). PANTHER version 11: expanded annotation data from Gene Ontology and Reactome pathways, and data analysis tool enhancements. Nucleic Acids Res 45, D183–D189.

Ong, J. Y. and Torres, J. Z. (2019). Dissecting the mechanisms of cell division. J Biol Chem 294, 11382–11390.

Reber, S. and Hyman, A. A. (2015). Emergent Properties of the Metaphase Spindle. Cold Spring Harb Perspect Biol 7, a015784.

Scheffner, M., Munger, K., Byrne, J. C. and Howley, P. M. (1991). The state of the p53 and retinoblastoma genes in human cervical carcinoma cell lines. Proc Natl Acad Sci U S A 88, 5523–7.

Schindelin, J., Arganda-Carreras, I., Frise, E., Kaynig, V., Longair, M., Pietzsch, T., Preibisch, S., Rueden, C., Saalfeld, S., Schmid, B. et al. (2012). Fiji: an open-source platform for biological-image analysis. Nat Methods 9, 676–82.

Sentein, P. (1977). Action of nocodazole on the mechanisms of segmentation mitosis. Cell Biol Int Rep 1, 503–9.

Wieser, S. and Pines, J. (2015). The biochemistry of mitosis. Cold Spring Harb Perspect Biol 7, a015776.

Yee, C., Krishnan-Hewlett, I., Baker, C. C., Schlegel, R. and Howley, P. M. (1985). Presence and expression of human papillomavirus sequences in human cervical carcinoma cell lines. Am J Pathol 119, 361–6.

